# Cellular landscape of adrenocortical carcinoma at single-nuclei resolution

**DOI:** 10.1101/2023.10.09.561514

**Authors:** David S. Tourigny, Barbara Altieri, Kerim A. Secener, Silviu Sbiera, Marc P. Schauer, Panagiota Arampatzi, Sabine Herterich, Sascha Sauer, Martin Fassnacht, Cristina L. Ronchi

## Abstract

Adrenocortical carcinoma (ACC) is a rare yet devastating tumour of the adrenal gland with a molecular pathology that remain incompletely understood. To gain novel insights into the cellular landscape of ACC, we generated single-nuclei RNA sequencing (snRNA-seq) data sets from twelve ACC tumour samples and analysed these alongside a previously published snRNA-seq data set from normal adrenal glands (NAGs). We find the ACC tumour microenvironment to be relatively devoid of immune cells compared to NAG tissues, consistent with known high tumour purity values for ACC as an immunologically “cold” tumour. Our analysis identifies three separate groups of ACC samples that are characterised by different relative compositions of adrenocortical cell types, including two populations (ACC 1 and ACC 2) that are specifically enriched in the most aggressive tumours and display hallmarks of the epithelial to mesenchymal transition (EMT) and dysregulated steroidogenesis, respectively. In addition to cell types associated with hypoxic and metabolic signatures (ACC 3 and ACC 4) prevalent among less-aggressive tumours, we also identified and validated a population of mitotically active adrenocortical cells (ACC M) strongly overexpressing genes *POLQ* and *DIAPH3* that possibly supports the expansion of malignant cell lineages. The smallest identified ACC specific cell type, ACC 5, displays characteristics of increased proliferation and growth factor signalling, and is therefore a potential progenitor-like or cell-of-origin candidate for the different lineages involved in adrenocortical carcinogenesis. Intriguingly, linage tracing suggests the fate adopted by malignant adrenocortical cells upon differentiation appears to be at least partly associated with the copy number or allelic balance state of the imprinted *DLK1*/*MEG3* genomic locus, which we verified by assessing DNA methylation status among samples from the three groups of tumours defined by their different cell type compositions. Our results therefore provide new insights into the cellular heterogeneity of ACC, indicating that genetic perturbations to a hierarchical cellular differentiation mechanism underlying healthy adrenocortical renewal and zonation may explain the molecular basis for disease pathogenesis.

## Introduction

Adrenocortical carcinoma (ACC) is a rare yet highly aggressive tumour of the adrenal gland (annual incidence 0.7-2 cases per million in USA; 5-year survival 50-60% early stage detection, 10-30% in later stages) with a molecular pathology that remains incompletely understood, which poses significant management challenges [1–5]. ACC is often associated with hypersecretion of various adrenocortical steroid hormones (usually cortisol and androgens) although can also be endocrinologically inactive meaning that clinical detection can be delayed until the later stages of disease [6]. ACC presents most frequently as sporadic, but can be associated with genetic syndromes in approximately 3-5% of cases [6,7]. Histological diagnosis of sporadic ACC is based on the modified Weiss scoring criteria, which includes parameters on tumour architecture, mitosis, and tumour invasion [8,9]. Immunohistochemistry (IHC) has proved an invaluable diagnostic and prognostic companion tool, and current practice is based on the use of markers including steroidogenic factor 1 (SF1) to recognise the adrenal origin of the tissue and the Ki-67 proliferation index to define the aggressiveness of the tumour [9–13].

Recent integrative [14] and pan-genomic [15] analyses have investigated the molecular attributes of sporadic ACC, with the latter contributing to The Cancer Genome Atlas ACC project (TCGA-ACC). These studies have highlighted a role for whole genome duplication and large-scale chromosomal copy number variations (CNVs), including near-ubiquitous overexpression of *IGF2* through loss of heterozygosity (LOH) or aberrant methylation of the maternal allele at the imprinted locus 11p15.5. Using Cluster of Cluster (COC) analysis of integrated genomic, DNA methylation, messenger RNA (mRNA) and micro-RNA (miRNA) expression data, three molecular subtypes of sporadic ACC were proposed to be consistent with grading based on histology and IHC [15]. Specifically, molecular subtype COC3 contained the most aggressive tumours, with COC1 showing the most favourable clinical outcome and COC2 presenting an intermediate prognosis.

Sporadic ACC is characterised by cancer initiating events that occur during renewal of post-developmental adrenocortical tissue [16,17]. The post-developmental human adrenal cortex is composed of three concentric zones of cells responsible for synthesising three different sets of steroid hormones: the outer zona glomerulosa (ZG, mineralocorticoids), zona fasciculata (ZF, glucocorticoids) and the inner zona reticularis (ZR, androgens). Renewal of this zonation is believed to involve at least one adult stem cell population sustained by a gradient of *WNT* signalling concentrated near the outer capsule, which becomes replaced by an increasing gradient of Protein Kinase A (PKA) signalling as cells are displaced to undergo centripetal migration and differentiation [18–20]. Exact mechanisms by which *WNT* and PKA signals interact to generate the different adrenocortical zonal cell types remain incompletely understood, but *WNT* modulators together with phosphodiesterase enzymes (PDEs) are suggested to achieve this balance by controlling intracellular levels of beta-catenin and cAMP, respectively [21–25]. Such a model is consistent with findings that mutations in *WNT* and PKA pathways are among the most commonly-occurring somatic events associated with various lesions of the adrenal cortex [2,14,15,26].

Although great progress has been made towards delineating the molecular mechanisms of adrenocortical development, renewal and pathogenesis in humans, very little remains known about the landscape of cell types involved in these processes. Notable exceptions include recent single-cell and single-nuclei studies on the foetal [27] and adult [28] human adrenal gland, respectively. While the latter also includes data from a cohort of adrenocortical adenoma (ACA) samples, equivalent data on ACC have hitherto been lacking. To better understand the composition of cell types and mechanisms involved in ACC pathogenesis, in the present study we assembled and analysed a cellular transcriptome atlas using a cohort of tissue samples from ACC patients.

## Results

### Cellular landscape of adrenocortical carcinomas

We conducted single-nuclei transcriptomic sequencing (snRNA-seq) on twelve tumour samples (six primary, three metastatic and three recurrent) from a total of eight patients (7F/1M, median age 58.5 yrs) with histologically confirmed diagnosis of ACC (Table 1 and Materials and Methods). Patients displayed a variety of steroid profiles at the time of resection, including four with mixed cortisol and androgens excess, one with cortisol excess alone, two with endocrine inactive tumours, and a single male patient with a rare, estrogen-secreting tumour. Following quality control and removal of genes mapping to mitochondrial and ribosomal genes from count matrices, normalisation and variance stabilisation was performed using Seurat [29]. Initial cell types were predicted for each sample individually using a merged snRNA-seq data set from six normal adrenal glands (NAGs) from a previous study as a reference [28], which are found to contain cells of adrenocortical origin together with adrenal medullary cells (AM), myeloid cells (MC), lymphoid cells (LC), fibroblast and connective tissue cells (FC), and vascular and endothelial cells (VEC). We then compared initial cell type predictions with annotations based on the same six cell types but informed by differential gene expression (DGE) analysis, performed following principal component analysis and unsupervised clustering after merging normalised count matrices for all ACC and NAG samples (Materials and Methods). Both sample-based and merged approaches to cell type inference confirm a relative excess of cells with adrenocortical origin and depletion of immune cell types in ACC compared to NAG samples (Figure S1), in line with reports of extremely high tumour purity and low immune and stromal infiltration in ACC [15,30,31].

**Table 1.** Demographic, clinical, and hormonal characteristics for the eight patients with adrenocortical carcinoma (ACC) and the six subjects with removed normal adrenal glands (NAG) included in the study. Abbreviations: ACC, adrenocortical carcinoma; M, male; F, female; N, not applicable; NAGE, normal adrenal gland deriving from endocrine inactive adenoma (EIA); NAGR, normal adrenal gland deriving from renal cell carcinoma (RCC); NGS, next generation sequencing. *Cases sequenced twice.

| Patient ID | Diagnosis | Sex | Age at surgery | ENSAT tumour stage at the time of diagnosis | Type of surgery | Hormone pattern at time of tissue sampling | DNA sequencing | Somatic variants in the analysed tissue (gene names) | Tumour size of the analysed tissue (mm) | Ki67% of the analysed tissue |
| --- | --- | --- | --- | --- | --- | --- | --- | --- | --- | --- |
| PACCp-1 | ACC | F | 36 | 3 | Primary | Cortisol & androgens | Targeted NGS | *TP53* | 170 | 20 |
| PACCp-2* | ACC | F | 72 | 1 | Primary | Cortisol & androgens | Targeted NGS | *CTNNB1* | 50 | 15 |
| PACCp-3 | ACC | F | 66 | 2 | Primary | Cortisol & androgens | Targeted NGS | *APC, NF1* | 82 | 10 |
| PACCp-4* | ACC | M | 71 | 1 | Primary | Estrogen | Targeted NGS | *KDM6A1, RB1, TP53* | 45 | 40 |
| PACCr-1* | ACC | F | 38 | unknown | Recurrence | Cortisol | Sanger seq | WT for *CTNNB1, GNAS* and *PKA* | 35 | NA |
| PACCr-2 | ACC | F | 61 | 2 | Recurrence | Endocrine inactive | Targeted NGS | none | 94 | 15 |
| PACCm-1* | ACC | F | 56 | 3 | Metastasis | Cortisol & androgens | Sanger seq | WT for *CTNNB1, GNAS* and *PKA* | 100 | 2 |
| PACCm-2 | ACC | F | 45 | 1 | Metastasis | Endocrine inactive | Targeted NGS | *KDR, MEN1, ZNRF3* | 25 | 50 |
| PNAGe-1 | NAG from EIA | F | 54 | NA | NA | NA | NA | NA | NA | NA |
| PNAGe-2* | NAG from EIA | F | 49 | NA | NA | NA | NA | NA | NA | NA |
| PNAGe-3 | NAG from EIA | M | 71 | NA | NA | NA | NA | NA | NA | NA |
| PNAGr-1 | NAG from RCC | M | 56 | NA | NA | NA | NA | NA | NA | NA |
| PNAGr-2* | NAG from RCC | M | 81 | NA | NA | NA | NA | NA | NA | NA |
| PNAGr-3 | NAG from RCC | M | 68 | NA | NA | NA | NA | NA | NA | NA |

Excluding cells with inconsistent annotations resulted in a final total of 39,364 cells for downstream analysis. We renormalised and remerged ACC and NAG count matrices for these cells, performing unsupervised clustering at higher resolution to reveal a total of 19 cellular clusters (Figure 1A). To identify the nature of cell types, DGE analysis was combined with a calculation of the log ratio of proportions (ROP) of ACC to NAG cells within each cluster (Figure 1B) (Materials and Methods). Using the rationale that a higher log ROP is indicative of a higher proportion of cells derived from cancer samples, we identified eight adrenocortical cell types that appear to be specific to ACC. Conversely, clusters with a lower log ROP include infiltrating cell types, medullary cells and two adrenocortical cell types identified with the ZG and ZR, based on expression of known markers such as *DACH1* and *SULT2A1*, respectively (Table S1) [28]. Four remaining adrenocortical cell types, which contained approximately equal proportions of cells from ACC and NAG samples (Figure 1B), are identified as representing a progressive continuum of cell types from the ZF based on expression of cytochrome P450 *CYP17A1* (ZF 1) and non-canonical *NOTCH* ligand *DLK1* (ZF 2 and 3) (number of ZF cells expressing *DLK1* in the normal adult adrenal gland has recently been shown to increase with patient age) [28,32], and a comparatively undifferentiated adrenocortical zone (UZ) expressing very few known marker genes (Table S1).

**Figure 1.**
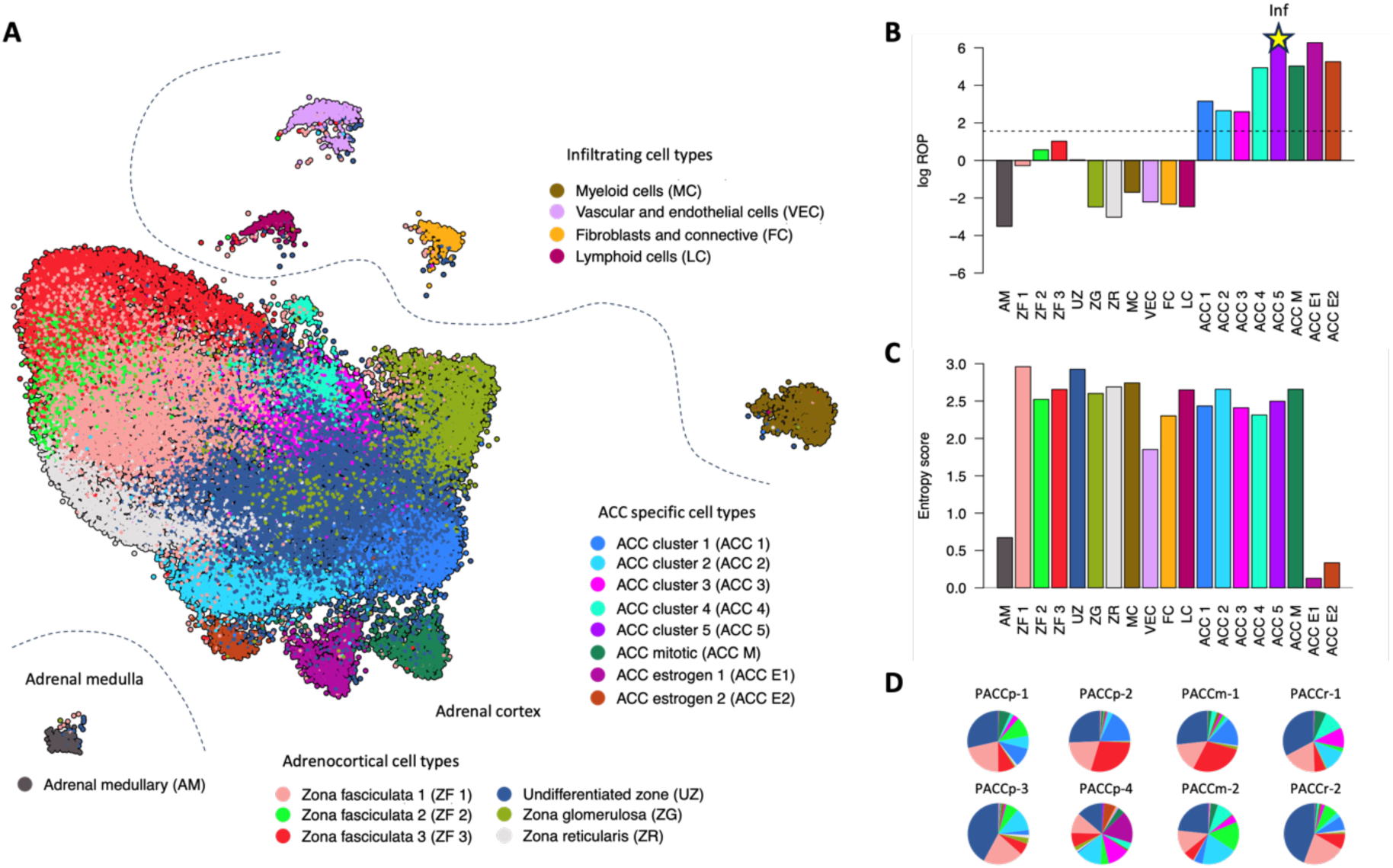
Landscape of cell types in ACC. (A) UMAP projection showing all cells in the merged data set of ACC and NAG samples, coloured by cell type. (B) Bar plot of log ROP scores (see Materials and Methods for details) for each cell type where the dotted line serves as a rough guide above which those that fall were denoted ACC specific. Note that ACC 5 consists entirely of cells from ACC samples meaning that its log ROP score is infinitely large. (C) Bar plot of patient-based Shannon entropy score (see Materials and Methods for details), indicating the degree to which each cell type is patient specific. (D) Pie charts showing a different representation of the distribution of cell types among the eight ACC patients included in this study (see Table 1 for details).

To quantify the distribution of cell types across ACC patients, we calculated a score based on Shannon entropy (Figure 1C) (Materials and Methods). Cell types that are highly specific to one patient have an entropy score close to zero, while the entropy score increases as cell clusters become distributed across patients. Two cell types with the lowest entropy score (ACC E1 and ACC E2) are found to consist almost exclusively of cells (99.3% and 97.7%, respectively) from PACCp-4 who was the only male patient in our cohort and had clinically excessive levels of circulating estrogen at the time of resection (Figure 1D). By comparison, the remaining six ACC specific cell types have higher entropy scores, reflecting the fact that they are represented in tumours from multiple patients in the study (Figure 1D).

### Characterisation of ACC specific cell types

Different ACC specific cell types were characterised using MSigDB hallmark gene set scores and identification of marker genes based on DGE analysis (Figure 2 and Tables S1 and S2) (Materials and Methods). ACC 1 cells score highest for *ANGIOGENESIS* but are most significantly enriched for the gene set *EPITHELIAL MESECHYMAL TRANSITION* (EMT) (adjusted p = 1.16e-82) and correspondingly their top marker genes (as ranked by p value) includes factors involved in cellular mechanics (*SPOCK1, COL11A1, CTNNA2*) and calcium signalling (*CALN1*, *CADPS, CACNA2D1, CACNB2*) (Figure 2B and Table S1), consistent with the role of calcium in regulating cytoskeletal and cell-matrix interactions during the EMT [33,34]. ACC 2, 3, and 4 cells were differentiated by their metabolic characteristics: ACC 2 cells score most significantly for the gene set *CHOLESTEROL HOMEOSTASIS* (adjusted p = 3.75e-203), consistent with top marker genes including those involved in cholesterol transport (*GRAMD1B*, *LDLR, SCARB1*), the mevalonate pathway (*HMGCS1*, *HMGCR*), sterol biosynthesis (*SQLE*, *FDPS*, *FDFT1, MSMO1*) and early steps in steroid hormone synthesis (*CYP11A1*, *FDX1*) (Figure 2B and Table S1). The top marker genes for ACC 3 are acyl-CoA synthetase *ACSM3* and *THUMPD1* (average log2 fold change = 1.36 and 0.71, respectively, adjusted p < 2.3e-308) (Figure 2B and Table S1), whereas ACC 4 scored highest for the gene sets *HYPOXIA* and *GLYCOLYSIS* (adjusted p = 6.11e-175 and 7.31e-108, respectively) (Figure 2A and Table S2). As displayed in Figure 2B, cells from ACC 4 correspondingly express the vascular endothelial growth factor *VEGFA* and glycolytic genes such as *PGK1* among other top marker genes (average log2 fold change = 0.47 and 0.45, adjusted p = 5.15e-171 and 5.68e-165, respectively) while the VEGR receptor gene *FLT1* is a top marker gene for the endothelial cell cluster VEC (average log2 fold change = 0.73, adjusted p < 2.3e-308). This suggests that ACC 4 represents a cell type that actively stimulates glycolysis and angiogenesis in response to hypoxic signals. ACC E1 and E2 cells specific to patient PACCp-4 score most significantly for the gene sets *ESTROGEN RESPONSE EARLY* and *EMT* (adjusted p = 3.49e-30 and 4.31e-27, respectively) and *G2M CHECKPOINT* and *CHOLESTEROL HOMEOSTASIS* (adjusted p = 6.32e-9 and 2.00e-08, respectively), respectively, and similarly share a subset of marker genes with the major ACC specific clusters ACC 1-4 (Figure 2B and Table S1).

**Figure 2.**
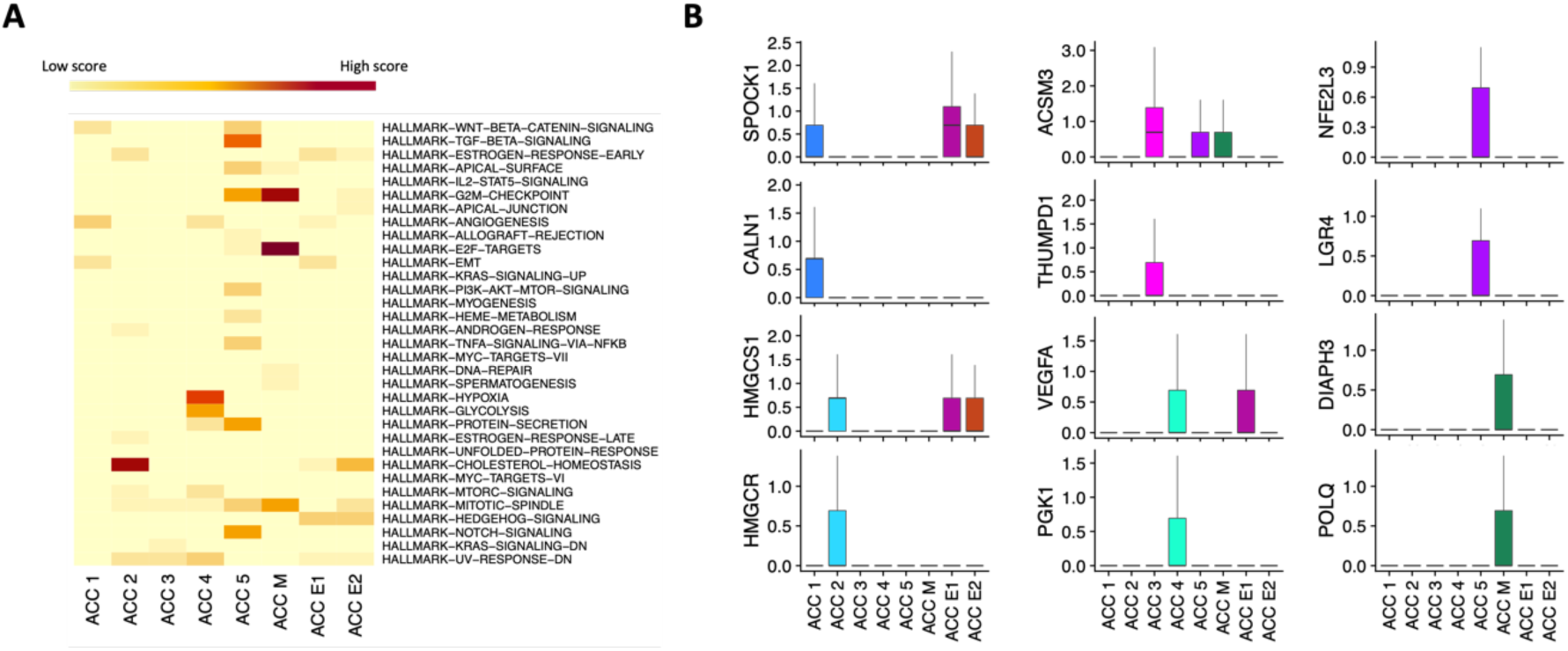
Characterisation of ACC specific cell types. (A) Heatmap of hallmark gene set scores for each ACC specific cell type (see Materials and Methods for details), coloured with a gradient from zero (lowest) to one (dark red). (B) Plots of selected top marker genes for each ACC specific cell type, where the vertical axis shows (SCT normalised) level of expression for the corresponding gene.

The two remaining ACC specific cell types, ACC 5 and ACC M, display hallmarks of proliferative and mitotic activity, respectively. ACC 5 is the cell type with by far the smallest representation in our data set (just 77 out of 39,364, less than 0.2 % cells) and its top markers include the marker of proliferation Ki-67 (*MKI67*) (average log2 fold change = 0.20, adjusted p = 4.5e-67) and components of the *WNT* (*LGR4*) and *NOTCH* signalling (*JAG1*) pathways (average log2 fold change = 0.29 and 0.25, adjusted p = 3.09e-08 and 0.0015, respectively), along with transcriptional factors such as *NFE2L3*, *SOX4* and the glucocorticoid receptor (*NR3C1*) (average log2 fold change = 0.36, 0.21 and 0.26, adjusted p = 3.86e-114, and 0.00016,respectively) that are important for determining proliferative cell fate in response to paracrine or endocrine signals (Figure 2B and Table S1) [35–39]. Correspondingly, ACC 5 cells score significantly for the gene sets *G2M CHECKPOINTS, TGF BETA SIGNALING*, *WNT BETA CATENIN-SIGNALING* and *NOTCH SIGNALING* (adjusted p = 4.29e-11, 2.02e-06, 0.007 and 0.009, respectively) (Figure 2A and Table S2). By comparison, ACC M contains almost ten-fold more representatives in our data set (734 out of 39,364, 1.9 % cells) and the top protein coding marker genes for this cell type are *DIAPH3* (diaphanous related formin 3) and *POLQ* (average log2 fold change = 0.46 and 0.35, respectively, adjusted p < 2.3e-308) as shown in Figure 2B and Table S1. *POLQ* is a DNA polymerase [40], and *DIAPH3* was recently reported to localise to the centrosome and regulate the assembly and bipolarity of the mitotic spindle during mitosis [41], which is consistent with indications that ACC M cells are actively engaged in mitotic activity. Correspondingly, ACC M scores most significantly enriched for the three gene sets *E2F_TARGETS* (adjusted p < 2.3e-308), *G2M_CHECKPOINTS* (adjusted p = 3.07e-213) and *MITOTIC_SPINDLE* (adjusted p = 3.23e-80) (Figure 2A and Table S2).

Since ACC M cells are found in relatively high numbers across all ACC samples while being almost entirely absent from NAG (high log ROP and entropy scores, Figures 1B and 1C), we investigated the possibility of using their top marker genes to distinguish ACC from normal or benign adrenocortical tissue at the bulk level. Both *POLQ* and *DIAPH3* were found to be significantly overexpressed in bulk RNA from ACC samples compared to NAG and ACA tissue, as reproduced in two different data sets (Figures 3A and S2) [42,43]. We next validated the expression of both *POLQ* and *DIAPH3* at the protein level using IHC and found that these proteins mark populations of cells that are found at significantly higher levels in ACC (n = 68) compared to NAG (n = 11) tissue (mean POLQ H-score ± Standard deviation (SD) = 135.6 ± 62.6 for ACC versus 73.5 ± 32.4 for NAG, respectively, p value = 0.0017; mean DIAPH3 H-score ± SD = 204.4 ± 50.9 for ACC versus 95.1 ± 25.6 for NAG, respectively, p value = 1.6e-06) (Figure 3B). Taken together, these results suggest that ACC M represents a population of mitotic adrenocortical cells that are generously and reproducibly enriched within malignant adrenocortical tumours.

**Figure 3.**
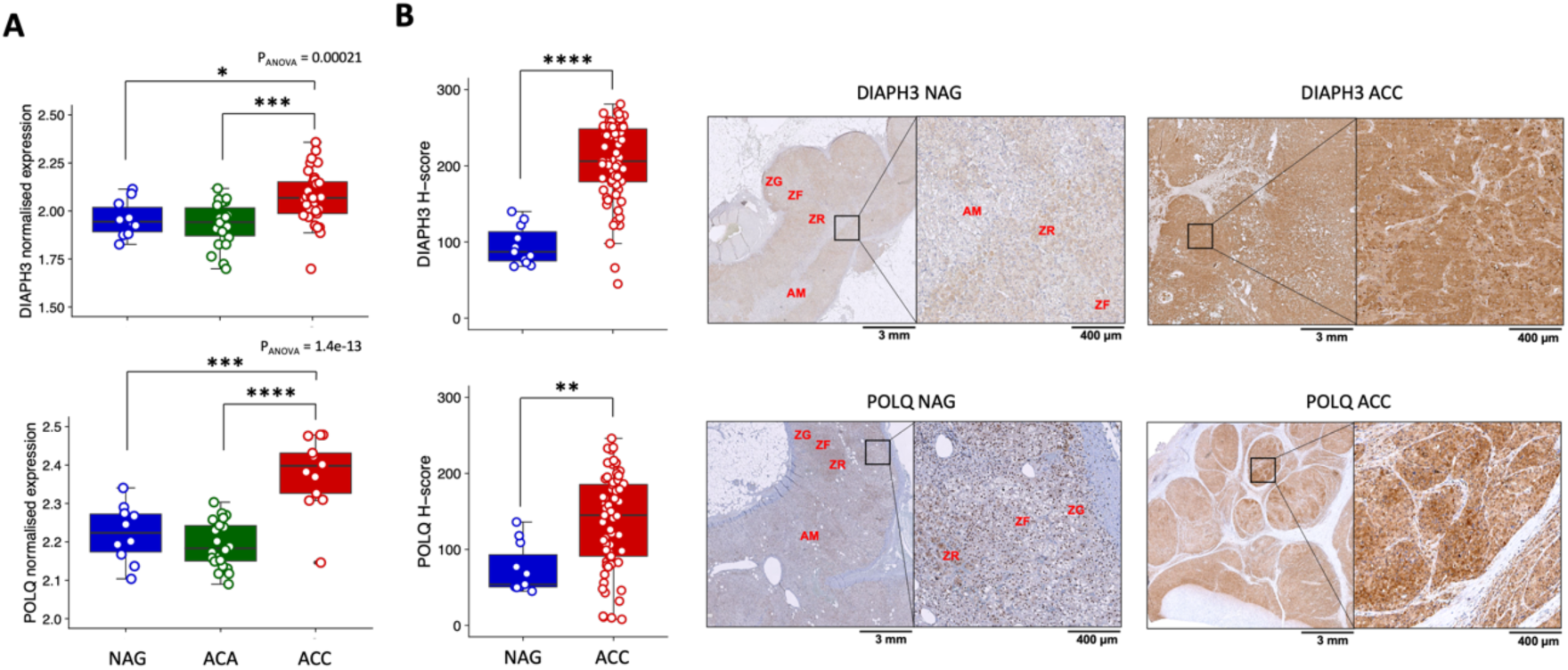
Validation of top marker genes for ACC M. (A) Summary of *POLQ* and *DIAPH3* expression in bulk RNA from Giordano et al., 2009 [43] with NAG (n = 10), ACA (n = 22) and ACC (n = 33) samples. (B) Validation of *POLQ* and *DIAPH3* expression at protein including representative images from IHC.

### Cellular compositions of molecular ACC subtypes

We next performed hierarchical clustering based on cell type composition, which separates out NAG samples and partitioned ACC samples into three distinct groups that mixed primary, recurrence and metastatic lesions (Figure 4A). Samples from group II are characterised by a significantly higher proportion of ACC 1 cells than groups I and III (p < 2.2e-16; 95% confidence interval (CI) [14.8%,16.3%] and [8.9%,10.9%] greater than groups I and III, respectively). Conversely, group I, which included samples from patient PACCp-4, is enriched for ACC 2 cells (p < 2.2e-16; 95% CI [10.6%,12.1%] and [5.5%,7.5%] greater than groups II and III, respectively) while group III has no significant difference in the proportions of ACC 1 and ACC 2 (p=0.45). Group I samples also have significantly higher proportions of the four remaining ACC specific cell types: ACC 3, 4, 5 and M (p < 2.2e-16; 95% CI [17.3%,19.2%] and [15.8%,17.9%] greater than groups II and III, respectively).

**Figure 4.**
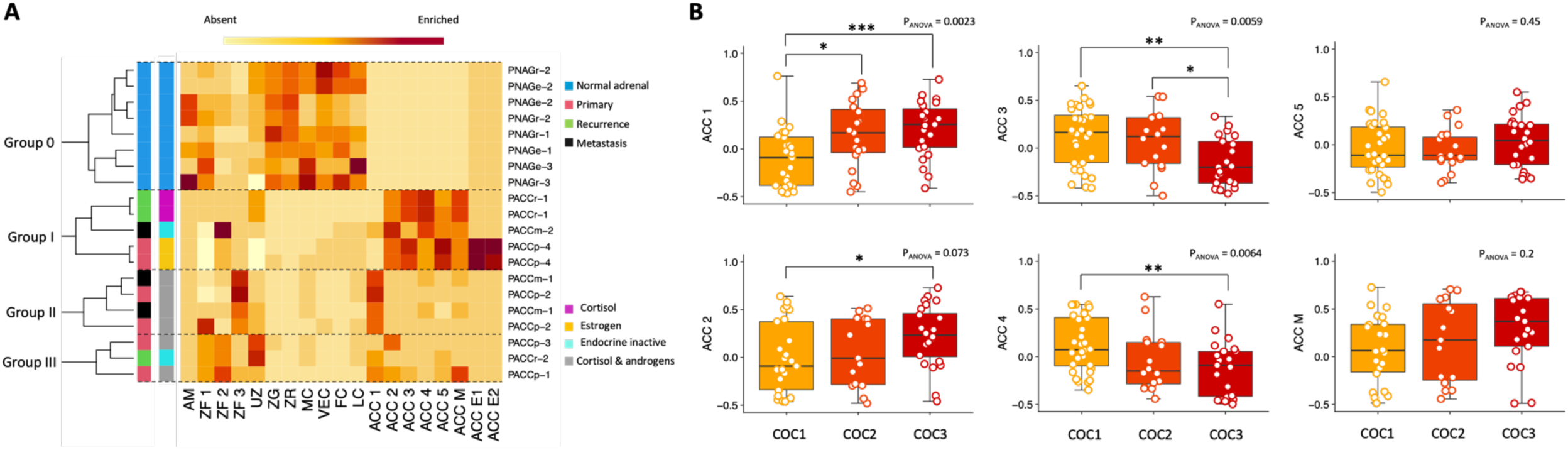
Grouping of ACC tumours by cellular compositions and molecular subtypes. (A) Heatmap showing the hierarchical clustering of NAG (Group 0) and ACC samples (Groups I, II and III) based on the relative proportions of cell types, coloured with a gradient going from zero (no cells of that type in corresponding sample, white) to one (all cells of that type in corresponding sample, dark red). (B) Scores for the six ACC specific (excludes ACC E1 and E2 from estrogen patient PACCp-4) single-cell signatures across TCGA-ACC samples grouped into their molecular subtypes.

To assess whether molecular ACC subtypes reflect associations with different cellular subpopulations, we generated single-cell signatures by taking the top 20 protein coding marker genes for each ACC specific cell type and used them to score TCGA-ACC samples based on gene set variation analysis (GSVA) [44] (Materials and Methods). COC3 samples scored significantly higher than COC1 samples for both ACC 1 and ACC 2 signatures (p value = 0.00053 and 0.021, respectively) as did COC2 for ACC 1 (p value = 0.019) (Figure 4B), suggesting these gene signatures are enriched in the more aggressive COC3 and intermediate COC2 ACC subtypes. By contrast, COC1 samples scored significantly higher than COC 3 samples for both ACC 3 and ACC 4 signatures (p value = 0.0019 and 0.0044, respectively) as did COC 2 for ACC 3 (p value = 0.027) (Figure 4B), suggesting these cell types are predominantly enriched in tumours with comparably good prognosis. No significant difference in scores for ACC 5 or ACC M were found across molecular subtypes. Thus, cellular populations ACC 1 and ACC 2, which are found to characterise group II and group I samples, respectively, are likely overrepresented in the most aggressive ACC molecular subtypes [15].

### Alternative adrenocortical cellular lineages in ACC

We noticed that a majority (68.3%) of cells from ACC samples were annotated as cell types UZ or ZF 1-3 (Figure 1D), and each have log ROP values close to zero implying that they are approximately equally represented in NAG and ACC samples (Figure 1B). This should be contrasted with the estimated average tumour purity value of 0.82 reported for ACC (higher than any other TCGA cancer type apart from chromophobe renal cell carcinoma) [15], which might therefore suggest that a large proportion of cells with a cancer genotype still adopt a normal phenotype. Reliable information on the genomic state of UZ and ZF cells is not accessible without additional DNA sequencing [45], but to gain insight into their nature in the context of other adrenocortical cell types we performed snRNA-seq-informed trajectory analysis using Slingshot [46] (Materials and Methods). Slingshot was chosen as the only method with a nearly perfect usability score that performs consistently across a wide range of evaluation criteria [47]; it is not fully reliable with disconnected regions in the dimensional reduction however, so we excluded cell types ACC M, ACC E1 and E2 from the trajectory analysis. Slingshot predicted six lineages based on a branched topology that contained branch points at UZ, ZF 1 and ACC 5 (Figure S3A). In addition to being enriched for a large number of signalling pathways and proliferative markers, ACC 5 also scored highest for an adapted version of the StemID score [48] (Material and Methods), consequently identifying ACC 5 as a candidate progenitor cell type for the linages obtained by Slingshot (Figure 5A).

**Figure 5.**
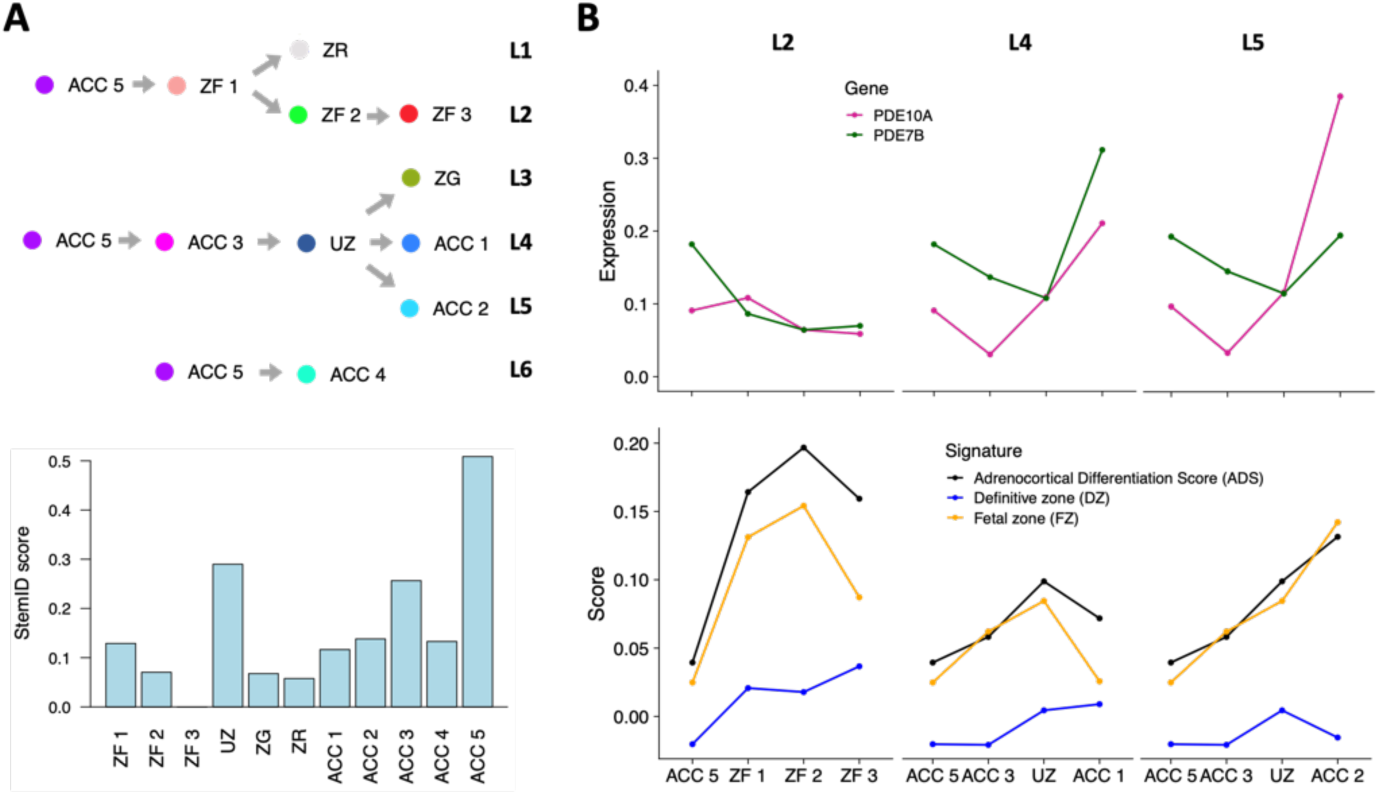
Adrenocortical lineages in NAG and ACC. (A) Graphical summary of adrenocortical cell types lineages obtained from trajectory inference with ACC 5 as a root node based on it having the highest StemID score ([48] and Materials and Methods) among all included cell types. (B) Expression levels (SCT normalised) across lineages L2, L4 and L5 for *PDE10A* and *PDE7B* (upper three graphs) and scores for published adrenocortical gene sets (lower three graphs).

Lineages L1, L2 and L3 are consistent with a centripetal migration and differentiation model of adrenocortical zonation where the ZF progressively matures (as measured by *DLK1* expression; average log2 fold change = 1.09 and 0.97, and adjusted p = 1.51e-153 and adjusted p < 1.51e-153 in ZF 2 and ZF 3, respectively) and the ZR emerges from the steroidogenic ZF phenotype (ZF 1), while the ZG emerges from a progenitor or intermediate cell type (UZ) [18–20]. Lineages L4 and L5 (terminating in ACC 1 and ACC 2, respectively) also pass through the UZ intermediate cell type and have sharply increasing expression of several PDE genes compared to lineage L2, where expression slightly decreases or remains approximately constant (Figure 5B). Increased PDE expression along lineages L4 and L5 may lead to a decrease in cAMP levels that blocks differentiation into the ZF or ZR phenotypes, which depends on PKA signalling in healthy adrenocortical tissue [22,23,49]. Figure 5B also displays L2, L4 and L5 lineage scores for three gene sets defined in previous adrenocortical studies: adrenal differentiation score (ADS) from the TCGA-ACC study [15], and the foetal zone (FZ, inner layer involved in early androgen biosynthesis similar to the adult ZR) and definitive zone (DZ, histologically similar to the postnatal ZG and outer ZF) signatures from a recent single-cell study of the developing adrenal gland [27]. The scoring patterns for ADS and FZ are closely related and ZF 2 and ACC 2 score higher than ZF 3 and ACC 1 for these signatures, respectively. Conversely, the scoring pattern is inverted for DZ (ZF 3 and ACC 1 score higher than ZF 2 and ACC 2 for this signature), and on further inspection we found that a significantly higher proportion of ZF 3 cells are found in group II samples (p < 2.2e-16; 95% CI [20.8%,22.9%] and [19.6%,22.0%] greater than groups I and III, respectively), as found previously for ACC 1. This highlights that group II samples are distinguished, not only by a greater proportion of ACC 1 cell types, but also by an increased proportion of cells that progress to mature a differentiation phenotype characterised by increased *DLK1* expression along the ZF lineage L2 (Figure S3B).

### Role of the imprinted DLK1/MEG3 genomic region in adrenocortical cell fate

Alongside imprinted gene *DLK1* that is suspected to play a role in healthy and diseased adrenocortical zonation [28,32], imprinted gene *IGF2* is another top marker for cell types ZF 2 and 3 at the ends of lineage L2 (average log2 fold change = 1.30 and 0.81, adjusted p = 6.52e-122 and adjusted p < 2.17e-302, respectively) (Table S1). The relevance of *IGF2* overexpression in ACC has been the subject of intense investigation [50–52], and both of these paternally expressed genes encode signalling ligands expressed within developing adrenal cortex with cognate receptors are found on mesenchymal cells within the adrenal capsule [27]. We also observed that *MEG3*, expressed from the maternal allele at the imprinted *DLK1/MEG3* chromosomal locus 14q32.2, serves as a top marker gene for the ACC 2 cell type (average log2 fold change = 0.77, adjusted p < 2.17e-302) from group I samples, but is not as strongly expressed by the ACC 1 and ZF 3 cell types enriched among group II samples (Figures S3C and S3D and Table S1). We therefore hypothesized that patterns of *DLK1*/*MEG3* expression across cell types (Figure S3D) may reflect different underlying genetic alterations at the imprinted 14q32.2 locus, which could potentially explain alternative cellular compositions of different ACC samples. To explore this, we used the top 20 marker gene single-cell signatures for ACC 1 and ACC 2 to score TCGA-ACC samples separated by chromosomal copy number state [53] of the 14q32.2 region. Strikingly, we found that samples that have undergone chromosomal gains in this region score significantly higher for the ACC 1 signature (p value = 0.046) while those that had undergone copy neutral LOH score significantly lower for the ACC 2 signature (p value = 0.01) (Figure 6A).

**Figure 6.**
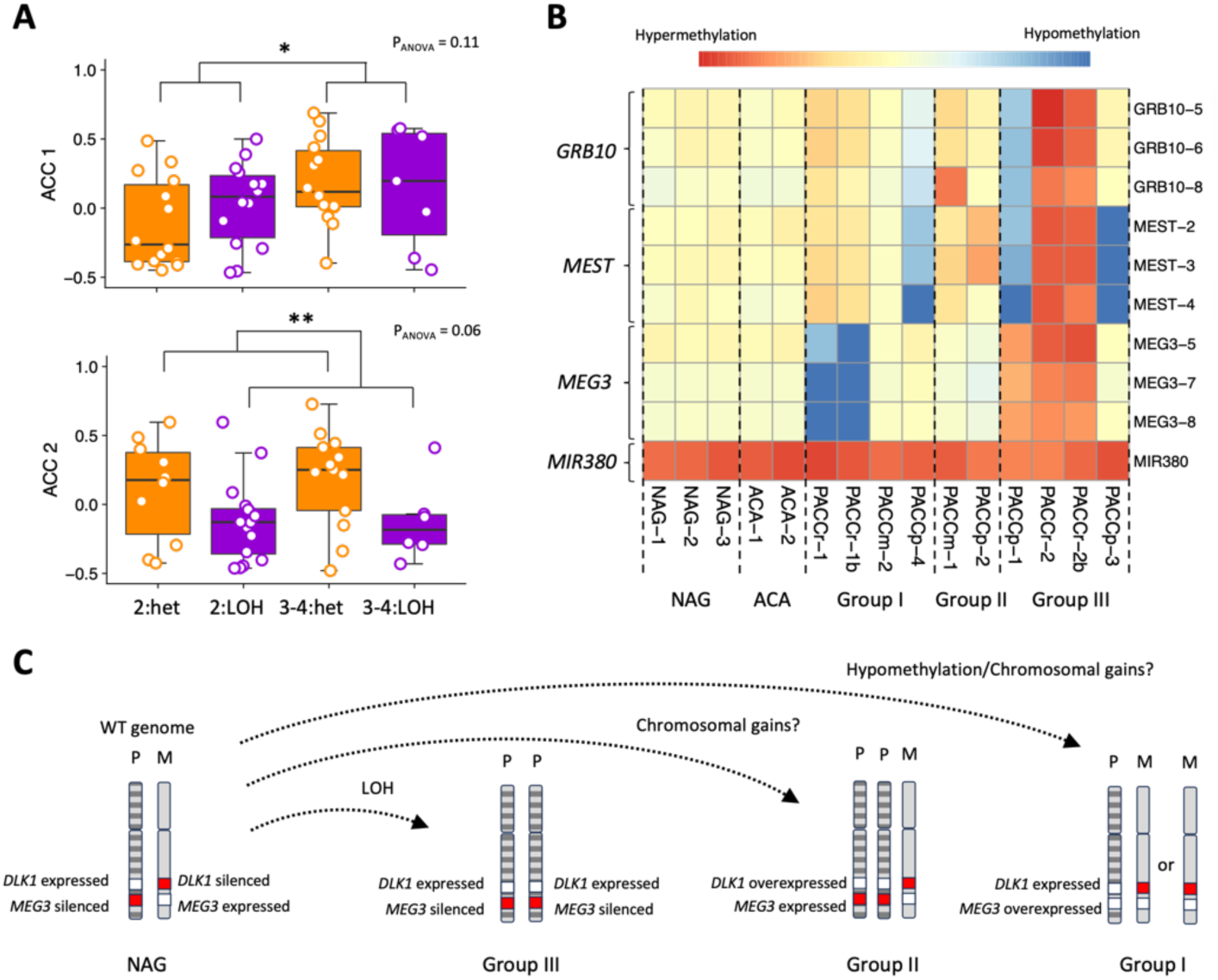
Associations of cell type compositions with state of imprinted 14q32.2 region. (A) Scores for single-cell signatures ACC 1 and ACC 2 across TCGA-ACC samples grouped by copy number state at the 14q32.2 region encoding *DLK1* and *MEG3*. Only copy number segments larger than 40Mbp are included and chromosomal gains of more than four are excluded. Heterozygous segments are those with A > 0, B > 0 (reflecting the counts for major allele A and minor allele B) while LOH are those with A > 0, B = 0: diploid wildtype or copy neutral LOH (2:het and 2:LOH, respectively) and chromosomal gains (3-4:het or 3-4:LOH). (B) Heat map showing the methylation status of probed genomic regions in NAG, ACA and ACC samples grouped according to cellular composition. (C) Cartoon schematic illustrating possible mechanisms for generating different ACC cellular compositions based on alterations at the imprinted 14q32.2 region. Group II samples, characterised by enrichment for ACC 1 and ZF 3 (*DLK1* high) populations, possibly associated with copy number gains of the parental (P) allele. Loss of methylation at the *MEG3* locus and/or copy number gains of the maternal (M) allele could underlie enrichment of ACC 2 (*MEG3* high) populations in group I samples, while both ACC 1 and ACC 2 cell types are comparatively depleted in group III samples associated with copy neutral LOH.

Finally, to corroborate the relationship between 14q32.2 chromosomal state and ACC cellular composition, we performed multiplex-ligation dependent probe amplification (MLPA) to assess the methylation status of this genomic region (Materials and Methods). In healthy cells, imprinting of 14q32.2 is controlled by methylation of the *MEG3* coding region on the paternal allele, which restricts its expression to the maternal (unmethylated) allele. We first confirmed that MLPA can reliably detect and distinguish hypo-, hyper- and heterozygous methylation states of *MEG3* (and known methylated regions *GRB10*, *MEST*, *MIR380*) using DNA isolated from paired patient blood samples to show these have the same methylation patterns as NAG and ACA controls (Figure S4). The assay was then repeated using DNA isolated from tumour samples from the eight ACC patients included in our study, revealing that two of the three tumours from group III (PACCp-1 and PACCr-2, confirmed by technical replicate) display hypermethylation of *MEG3* associated with complete loss of expression of this gene from the maternal allele (Figure 6B). Hypermethylation and silencing of *MEG3* expression can arise via several different mechanisms (including LOH of the maternal allele, Figure 6C) that are not possible to distinguish using MLPA. However, finding this alteration to occur only within samples from group III is entirely consistent with the characterisation of this group by a relative absence of ACC 1 and ACC 2 cellular populations, which are potentially associated with retention or additional gains of the maternal allele (Figure 6C). Furthermore, *MEG3* methylation is found to be entirely absent from one tumour in group I (patient PACCr-1, confirmed by technical replicate) (Figure 6B), implying that loss of imprinting at the paternal allele could be one possible mechanism driving *MEG3* overexpression within the ACC 2 cell type from group I samples (Figure 6C). Taken together, these results strongly suggest that the copy number state and balance of alleles at the *DLK1*/*MEG3* 14q32.2 locus are associated with the relative cellular compositions of different ACC tumours.

## Discussion

Our understanding of ACC pathogenesis has increased considerably over the past decade following in-depth characterisations of the molecular processes involved in development, homeostasis and disease of the adult adrenal gland [2,3,14,15,19–21,27,28]. In the present study, we contribute to this emerging field by providing a cellular transcriptome atlas of ACC at single-nuclei resolution that complements existing data and reveals a mixture of cell types present within the malignant adrenal cortex. Based on a detailed comparison with tissue from healthy adrenal gland, we find a low amount of immune infiltration confirming that ACC is a “cold” immune infiltrate tumour and consistent with previous reports of high tumour purity values [15,30,31]. We identify at least eight cell types that are specific to ACC, with at least two that are highly specific for the rare case of a male patient with excess levels of estrogen. ACC samples from our patient cohort separate into three predominant groups (that mix primary, metastatic, and recurrent tumours) according to their underlying composition of the six remaining ACC specific cell types (ACC 1-5 and ACC M).

The ACC sample group II is enriched for cell type ACC 1 that displays hallmarks of the EMT pathway as mediated by cytoskeletal and cell-matrix remodelling driven by calcium signalling [34] in addition to having a greater proportion of age-associated, *DLK1* and *IGF2* expressing ZF cells (cell type ZF 3). Conversely, the ACC sample group I is enriched for cell type ACC 2 that displays hallmarks of increased cholesterol uptake, early steroid metabolism and *MEG3* expression. Cell type ACC 2 may therefore account for aberrant steroidogenesis in a subgroup of ACC patients, including those that are deemed clinically endocrine inactive but still may secrete large quantities of steroid hormone precursors [54]. ACC samples from group III contained far fewer proportions of ACC 1 and 2 cell types, both of which were found to have signatures that are enriched in more aggressive COC3 and COC2 tumour subtypes within the TCGA-ACC [15]. Group III samples could therefore represent ACC tumours with comparably better outcome since they appear to lack these cell types. ACC 3 and ACC 4 cell type signatures, found mainly in group I samples, display hallmarks of increased fatty acid metabolism and hypoxia/glycolysis (ACC 3 and 4, respectively) that were also preferentially enriched in less-aggressive, COC1 tumours. One possible explanation for this observation is that metabolic perturbations blocking adrenocortical steroidogenesis can inhibit malignant ACC progression [55].

Consistent with existing knowledge on the increased expression of cell division and proliferation markers in ACC [13], two remaining ACC specific populations ACC M and ACC 5 both display hallmarks of increased DNA replication and mitotic activity. The more abundant cell type, ACC M, is characterised by the top marker genes *DIAPH3* and *POLQ* that we verified are also overexpressed at the protein levels in ACC. *POLQ* is a known DNA polymerase [40] and *DIAPH3* is a relatively uncharacterised protein that has recently been shown to localise to the centrosome during mitosis [41], and proposed as a biomarker for prostate and colorectal cancer [56,57]. Of note, both *POLQ* and *DIAPH3* are among the top marker genes for transit amplifying cells (TACs) in a recent single-cell data set comparing human colorectal polyps with normal tissue [58], suggesting that ACC M might represent a mitotic phenotype with increased prevalence in the diseased state of many self-renewing tissues such as the colon, intestine and adrenal cortex [59]. In this way, increases in numbers of cells from the ACC M population could drive an enrichment of malignant ACC 1 and ACC 2 cell types that are associated with more aggressive ACC tumours.

ACC 5 was identified as the least abundant ACC specific cell type and is characterised by high scores for *WNT*, *TGF*-beta and *NOTCH* signalling pathways and top marker genes *MIK67*, *LGR4, JAG1, SOX4, NFE2L3* and *NR3C1*. Among these top marker genes, expression of the *RSPO3* receptor *LGR4* was also recently detected in the developing adrenal cortex [27] and its mutation is known to drive aberrant adrenocortical zonation [60]. *RSPO3* itself is a *WNT* signalling coactivator that is proposed to play a key role in maintaining the adrenocortical stem cell niche near the adrenal capsule [20,21,24,25,27,28]. This model is once again analogous to the mechanism for maintenance of the *LGR5*-positive (homologue *RSPO3* receptor) stem cell compartment in colon or intestine, whose perturbation during carcinogenesis involves aberrant differentiation of TACs into absorptive or secretory cell types based on levels of *NOTCH* signalling [59,61–63]. In further support of this model, we have demonstrated that adrenocortical lineages and cellular compositions of the different ACC sample groups are associated with abnormal copy number states and allelic imbalances of the imprinted 14q32.2 locus, which encodes the non-canonical *NOTCH* ligand *DLK1* and maternally expressed gene *MEG3*. Silencing of *MEG3* through LOH of the maternal allele could reduce the rate of differentiation into cell types ACC 1 and ACC 2, which are underrepresented among group III samples and instead associated with more aggressive COC2 and COC3 tumours. This is consistent with previous results that show LOH of the maternal *DLK1*/*MEG3* allele is ubiquitous across a subset of ACC tumours with comparably favourable outcome [14,64]. While loss of *MEG3* expression may provide a selective advantage in early stages of disease [65–67], this alteration perhaps prevents progression into more advanced stages of ACC.

These data have several limitations, including a relatively small patient cohort due to the restricted availability of biological tissue for such a rare disease and the fact that snRNA-seq as a method can have limited sensitivity at low gene expression levels. The ability to expand single-cell profiling to larger patient cohorts and include more sequencing reads per cell will help to further advance our initial findings and hypotheses. Although knowledge of the molecular sequence of events involved in the pathogenesis of ACC remains far from complete, the present study has provided some novel insights into the vast landscape of cells and genes that are involved in this process. Our results therefore constitute an essential contribution to the biological and clinical comprehension of this devastating disease.

## Materials and Methods

### Patients and clinical details

Demographic, clinical and histopathological data for the eight patients with ACC included in the present study were collected from patients’ records and are summarised in Table 1. Diagnosis of ACC was histologically confirmed according to the current European guidelines [6]. Baseline ENSAT tumour stage, as well as Weiss score and ki67 proliferation index of the analysed tissues were collected. Steroid hormone levels were measured at the time of surgery (i.e., at resection of tumour tissue evaluated in the present study) using commercially available analytical procedures as previously reported [68]. Table 1 also includes the demographic characteristics of six subject with NAG deriving from the tissue surrounding EIA or from adrenalectomies performed during surgery for renal cell carcinoma, which were included in a previous study on snRNAseq [28] and used as reference. The study was approved by the ethics committee of the University of Würzburg (No. 93/02 and 88/11) and written informed consent was obtained from all subjects.

### Sample selection, preparation, and snRNA sequencing

Nuclei were isolated from snap-frozen samples derived from ACC tissue dissected by an expert pathologist and obtained during tumour resection using the protocol previously described in [28,69]. Quality and purity of isolated nuclei were confirmed by microscopy and yield was quantified using a Neubauer chamber. Salinization, reverse transcription (RT) and library preparation were performed according to the inDrop^TM^ system from 1CellBio [70] on chips with hydrogel beads, with library preparation following the CEL-Seq2 protocol [71] where the RT product was first digested by ExoI and HinFI and purified using AMpure XP beads (Beckman Coulter, USA). cDNA was quantified by qPCR (Roche Light Cycler 480 Instrument II, Switzerland) and pooled for sequencing using an Illumina HiSeq 2000 intrument with 60 bases for read 1, 6 for the Illumina index, and 50 for read 2.

### snRNA-seq data processing and analysis

FASTQ files containing raw snRNA-seq reads were generated using the Illumina bcl2fastq tool and processed with the zUMI pipeline (version 2.5 – default parameters) [72]. Count matrices were then used as input into Seurat (version 4.2) [29], which was used to create objects for all downstream processing and analysis in the *R* programming language. Initial quality control included filtering out cells that contained less than 100 or more than 2500 features, or more than 4000 total counts, after removing genes that mapped to ribosomal and mitochondrial coding or pseudo coding regions. We also repeated the analyses without removing these genes to confirm that this step did not affect the scientific conclusions of this paper.

Normalisation was performed on a sample-by-samples basis using the Seurat function SCTransform(), and initial cell types for ACC samples were predicted the merged data set of NAG samples as a reference with the Seurat functions FindTransferAnchors() and TransferData(), as described by the Seurat vignettes available online at that time. Following initial cell type predictions, normalised count matrices for ACC and NAG samples were merged and unsupervised clustering (resolution = 0.1) was performed using the top 30 principal components. Differential gene expression analysis performed on merged data using the Seurat function FindAllMarkers() provided a second route to initial cell type identification, and cells without a consistent annotation in both methods were excluded from downstream analysis. The filtered count matrices obtained after excluding these cells were then renormalised and remerged to create a final count matrix that was used for unsupervised clustering (resolution = 0.6) and final annotation of cell types based on differential gene expression and hallmark gene set analysis.

Gene set scores for cell types were calculated based on a customised version of single-cell gene set analysis by adapting the Seurat functions AddModuleScore() and FindAllMarkers() as previously described in [73]. Log ratio of proportions (log ROP) scores were obtained by dividing by the total number of cells in either sample type (ACC or NAG) and then taking the natural logarithm of the ratio of these proportions within each cell type. To calculate the Shannon entropy of each cell type, proportions were instead calculated on a patient basis after removing NAG samples and the function Entropy() from the *R* package DescTools was applied. Slingshot [46] was used according to the authors instructions by applying the functions getLineages() and SlingshotDataSet() with default settings for trajectory inference. The version of the StemID score used in this study is equivalent to that described in [48] where the estimated transcriptome entropy (with lowest entropy subtracted) for each cell type is multiplied by the number of links extending from that cell type as reported by Slingshot.

### Immunohistochemistry

Formalin-fixed paraffin-embedded (FFPE) slides of 79 adrenal tissues, including 68 ACC and 11 NAG were used to validate the expression of DIAPH3 and POLQ at protein levels by immunohistochemistry (IHC). Clinical data and pathological characteristics of the evaluated tissues are reported in Table S3. IHC was performed as previously reported [28]. Briefly, FFPE sections were deparaffinized and rehydrated in descending graded series of ethanol and antigen retrieval was obtained in citric acid monohydrate buffer (pH 6.5) at high temperature. 20% human AB serum was used for blocking of unspecific binding. Primary antibodies (anti-DIAPH3, #14341-1-AP, from Proteintech and anti-POL, #ab111218, from AbCam) were incubated 1 hour at room temperature. As negative control, N-Universal Negative Control Anti-Rabbit or Anti-Mouse (IS600 and IS750, respectively, Dako, Glostrup, Denmark) was used. Signal amplification was obtained by HiDef Detection HRP Polymer System (954D-50, Medac Diagnostika, Germany) followed by DAB substrate kit (957D-30, Cell Marque, USA). Finally, slides were incubated 2 min with Mayer’s haematoxylin (T865.1, Carl Roth, Germany) for the nuclei counterstaining. Images of the stained tissues were acquired with Leica Aperio Versa Brightfield scanning microscope (Leica, Germany) using same setting parameters to avoid biased information. The evaluation of the staining was performed by the automated image software analysis (Aperio ImageScope).

### Methylation status multiplex-ligation dependent probe amplification

Methylation status multiplex-ligation dependent probe amplification (MS-MLPA) was performed according to the manufacturer’s protocol using the SALSA MS-MLPA Probemix ME032-B1 UPD7-UPD14-v03. The probemix contains 14 reference probes and 32 specific probes for the regions 7p12.2, 7q32.2, 14q32.2 and 14q32.31, including the *GRB10*, *MEST*, *DLK1*, *MEG3* and *RTL1* genes. Ten of these specific probes contain a *Hha*I recognition site and provide information about the methylation status. 150 ng of DNA were used in each MS-MLPA reaction.

Hybridization, ligation and PCR were performed in a Biometra 96-well PCR thermal cycler. Fragment separation was done by capillary electrophoresis on a CEQ8000 capillary sequencer (ABSciex). Peaks are size-called and assigned to MLPA-probes by GenomLab GeXP-Fragment Analysis software. Quality check covers DNA concentration, MLPA-reaction, DNA-denaturation and *HhaI*-digestion. Peak height in relative fluorescence units (rfu) was used for quantification. After normalization, dosage quotients were calculated and compared between amplification products of the digested versus the undigested sample.

### Statistical analysis

All statistical tests reported in the main text were performed in the *R* programming language. Adjusted p values for marker genes and gene set scores were obtained using the Seurat function FindAllMarkers (), which uses a Wilcoxon signed-rank test with Bonferroni adjustment. The prop.test() function (default values) was used to test the null hypothesis that proportions of cell populations in two groups of samples are the same, with a two-sided alternative distribution and confidence level of 95%. GSVA [44] was used to score TCGA-ACC samples by cell type signatures derived from marker gene sets. The comparison of means of multiple sample groups was performed using an ANOVA test implemented in the function stat_compare_means() and p values for comparisons of pairs of groups were calculated using Wilcox method. Absolute p values for paired comparisons are reported in the main text while the annotations “*”, “**”, “***” and “****” have been used to denote p values less than 0.05, 0.01, 0.001 and 0.0001 in figures, respectively.

## Data and code availability

Raw sequencing data associated with this study have been uploaded to NIH Sequence Read Archive (SRA) under BioProject accession number PRJNA1024912 accessible via the link http://www.ncbi.nlm.nih.gov/bioproject/1024912. All code for reproducing the analyses presented in this article are freely available at https://gitlab.com/davidtourigny/adrenal-cortical-carcinoma.

## Acknowledgements

The authors are grateful to Ms. Michaela Haaf for coordinating the clinical data entry in Würzburg of the ENSAT registry. This work has been carried out with the help of the Interdisciplinary Bank of Biomaterials and Data of the University Hospital of Würzburg and the Julius Maximilian University of Würzburg (IBDW). The implementation of the IBDW has been supported by the Federal Ministry for Education and Research (grant number FKZ: 01EY1102).

## Funding

This work has been supported by the Deutsche Forschungsgemeinschaft (DFG) (project FA-466/8-1, RO-5435/3-1 and 405560224 (M.F., C.L.R., and S.S.)) and within the CRC/Transregio (project number: 314061271 – TRR 205), and the Deutsche Krebshilfe (70113526 M.F. and C.L.R.).

## Supplementary Material

### Legend to Supplementary Tables

**Table S1.** Marker genes for the 19 cell types reported in this study.

**Table S2.** Hallmark gene set scores for the 19 cell types reported in this study.

**Table S3.** List of samples included in marker validation by IHC.

### Legend to Supplementary Figures

**Figure S1.**
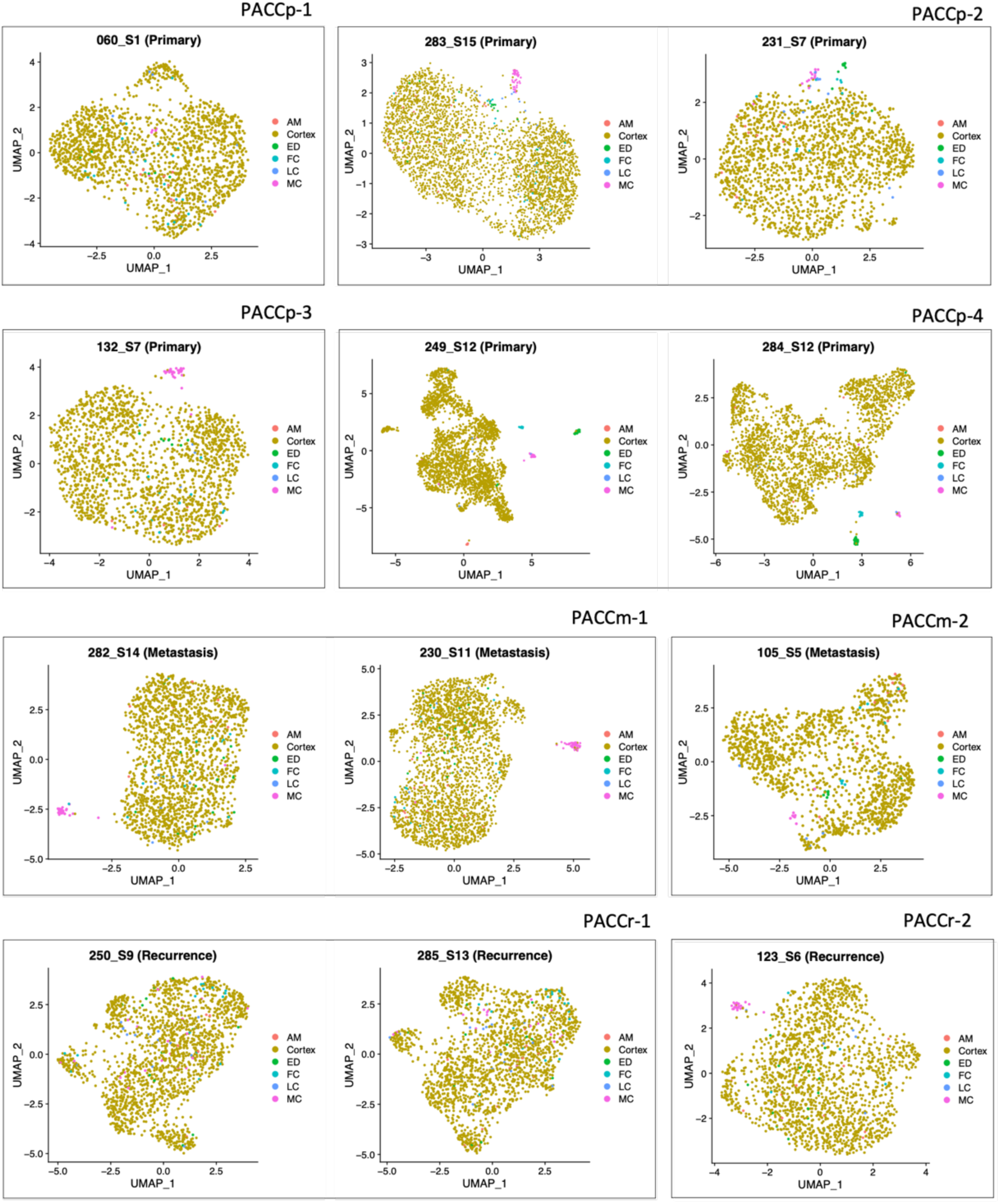
Initial cell type predictions on an individual sample basis using a merged data set of NAG samples as a reference. UMAP projections show all cells for each ACC sample, coloured by predicted cell type. Patients PACCp-2, PACCp-4, PACCr-1 and PACCm-1 were sequenced twice, as indicated in Table 1.

**Figure S2.**
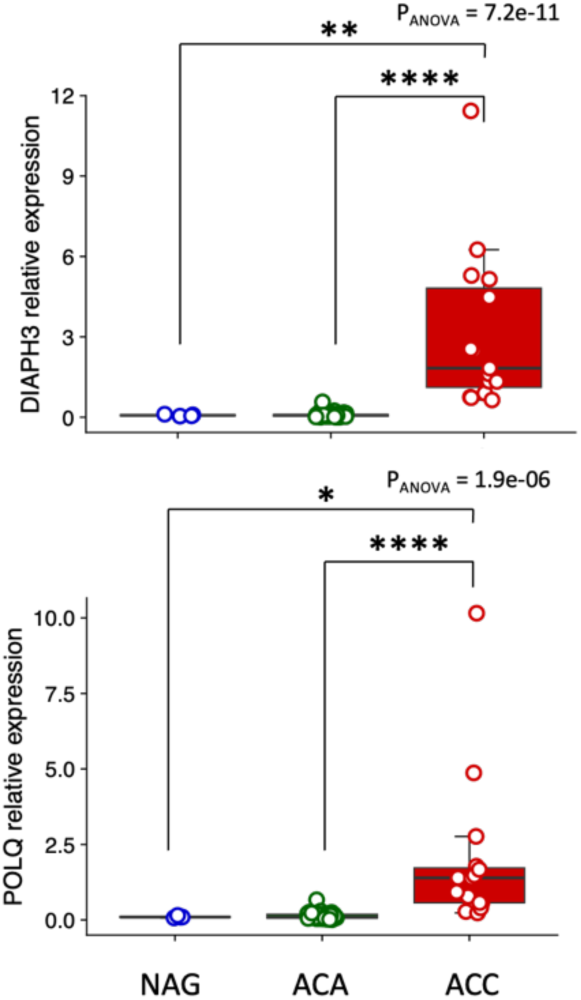
Summary of *POLQ* and *DIAPH3* expression in bulk RNA from Di Dalmazi, Altieri et al., 2012 [42] with NAG (n = 4), ACA (n = 57) and ACC (n = 15) samples.

**Figure S3.**
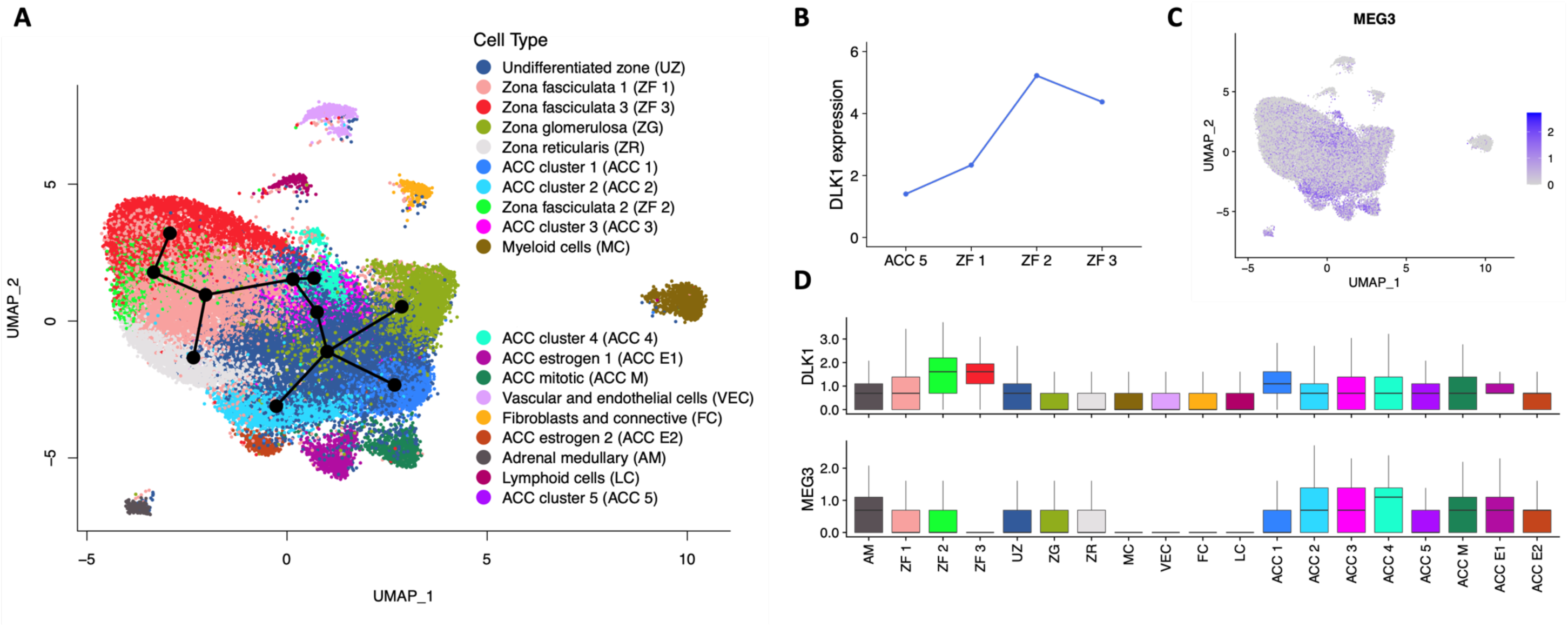
(A) UMAP projection overlayed with the branched trajectories obtained from Slingshot. (B) Average expression levels for *DLK1* along the lineage L2. (C) UMAP projection showing *MEG3* expression levels and (D) expression levels of *DLK1* and *MEG3* across cell types.

**Figure S4.**
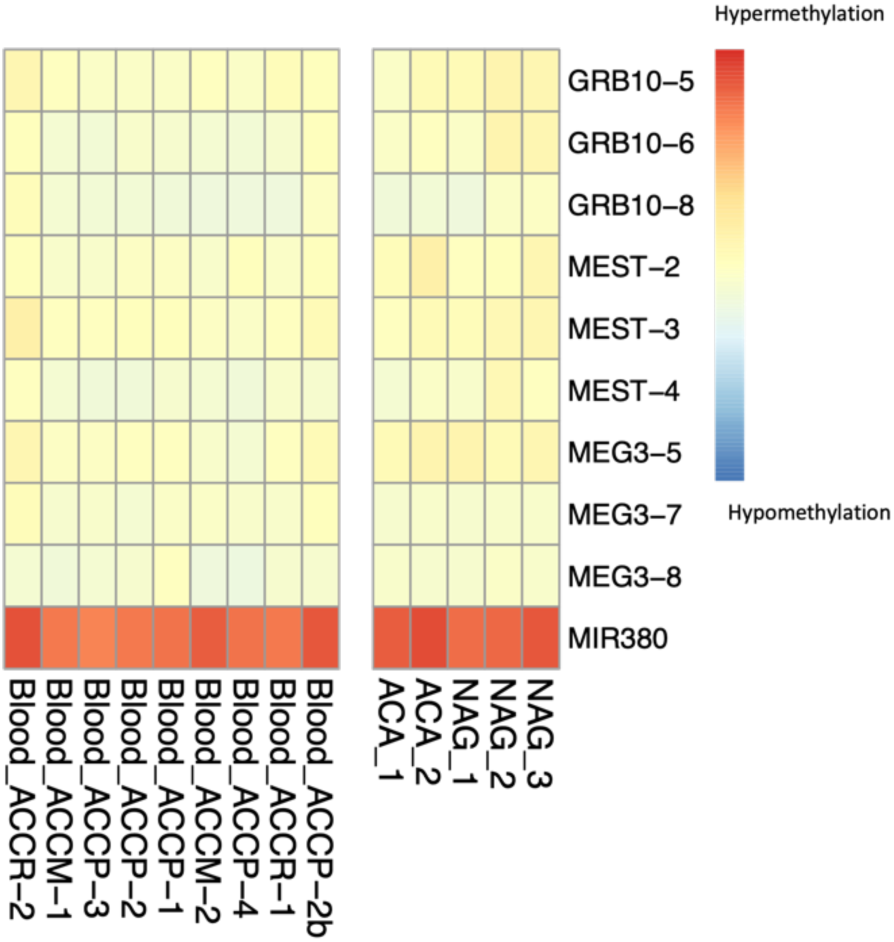
Heat map showing the methylation status of probed genomic regions in blood samples paired with tumour samples, and control NAG and ACA samples.

## Notes

https://gitlab.com/davidtourigny/adrenal-cortical-carcinoma

http://www.ncbi.nlm.nih.gov/bioproject/1024912

### Competing Interest Statement

The authors have declared no competing interest.

