## Supplementary material for "Cellular landscape of adrenocortical carcinoma at single-nuclei resolution": Table S3

| Parameter | NAG | ACC |
| --- | --- | --- |
| Number of evaluated tissues | 11 | 68 |
| Sex:  Female  Male | 5 (45.4)  6 (54.6) | 43 (63.2)  25 (36.8) |
| Age, years | 65 (59-76) | 49 (41-59) |
| Tissue size, cm | 3.9 (2.9-5.0) | 9.0 (6.0-14.0) |
| Tumor localization:  Primary  Local recurrence  Distant metastasis | n.a. | 59 (86.8)  5 (7.4)  4 (5.9) |
| Hormone secretion:  Inactive  Glucocorticoid (alone or with other steroids)  Androgens  Aldosterone  Unknown | n.a. | 15 (22.1)  25 (36.8)  6 (8.8)  1 (1.5)  21 (30.9) |
| ENSAT tumor stage:  I-II  III  IV | n.a. | 31 (45.6)  22 (32.4)  15 (22.1) |
| Resection status at primary diagnosis:  R0  R1-R2  RX  Unknown | n.a. | 43 (63.2)  16 (23.5)  6 (8.8)  3 (4.4) |
| Weiss score | n.a. | 6 (5-8) |
| Ki67% | n.a. | 20 (10-20) |

Continuous variables were reported as median (lower quartile - upper quartile), whereas categorical variables as number (percentage). Abbreviation: ENSAT, European Network for the Study of Adrenal Tumors; n.a.: data not applicable; R0, complete tumor resection; R1-R2, microscopically (R1) or macroscopically (R2) not completely resected tumor; RX, resection status not assessed.
